# Deep Learning Driven Investigation of Nanoplastic Impacts on Soil Protist Behavior in Soil Chips

**DOI:** 10.1101/2025.09.04.674150

**Authors:** Hanbang Zou, Wei Ying, Paola Micaela Mafla Endara, Fredrik Klinghammer, Jingmo Bai, Hanwen Kang, Edith C. Hammer

## Abstract

Nanoplastics are emerging contaminants that have a significant impact on soil microorganisms. To fully understand the effect of plastic contamination on soil ecosystems, it is necessary to advance techniques that can monitor nanoplastic-microbe interactions under realistic conditions. In this work, we investigated the effects of nanoplastic contamination on a community of soil protists monitored through microfluidic soil chips, and analysed changes in their behavior via microscopy videos and a deep learning approach. The presented method employs a deep learning-based detection model combined with a transformer-based matching model for video frame interpolation, enabling accurate reconstruction of protist movement trajectories and velocities within soil chips. The results revealed reduced movement velocities for the groups of flagellates and ciliates under high nanoplastic conditions, a 24-30% reduction at a marginal significance level, while amoebae were unaffected. Our trajectory data provides novel insights into how protists navigate soil-like structures. By facilitating comprehensive assessments of protist–environment interactions, it opens new avenues for understanding their ecological roles and the broader implications of hazardous contaminants in both soil and aquatic ecosystems at microbial community level without need for culture extraction. This proof-of-concept system enables continuous, high-throughput monitoring of soil protist behavior and can be readily adapted to investigate protist responses to diverse chemical and physical soil hazards.

## 1. Introduction

Plastics are synthetic or semi-synthetic materials composed of carbon-based polymers that do not occur naturally. These polymers are typically derived from fossil fuel sources such as crude oil and natural gas and are combined with various chemical additives to enhance their properties, including durability, flexibility, and resistance to environmental factors [1]. Unfortunately, the environmental impact of plastics is not fully understood. Most plastic materials are either non-biodegradable or degrade at an exceptionally slow rate, often persisting in the environment for decades or even centuries [2]. This persistence results in the accumulation of plastic waste in terrestrial and aquatic ecosystems, posing challenges to waste management systems and contributing to pollution on a global scale. Micro- and nanoplastics (MNPs) were first recognized as potentially hazardous pollutants to ecosystems in 2004 [3]. Since then, awareness of the risks posed by MNPs has grown significantly [4], but research remains limited in soil ecosystem [5, 6]. Most research on the effects of microplastics has focused on changes in microbial community structure and composition. However, the knowledge about ecophysiology and ecosystem functions we have gained remains limited, constrained by the tools and methodologies currently available. Soil protists, broadly defined as single-celled eukaryotes [7], contribute to approximately twice the global biomass of animals [8] and play vital roles as bacterial predators, facilitating the transfer of carbon from prokaryotic biomass to higher trophic levels while significantly influencing bacterial populations [9]. Protists can move over long distances in presence of continuous water films in soil [10], up to several centimeters per day [11]. In saturated conditions with wide pore spaces, some species can reach speeds of 300-500 *µ*m/s, whereas in complex and narrow pore spaces their movement slows to as low as 20 *µ*m/s [12]. They use different locomotion strategies to move within pore spaces and are broadly separated into the functional groups of flagellates, ciliates, and amoeba. In a generalized sense, flagellates move through a whipping motion of their one or more flagella while ciliates move through concerted movement of their many cilia, and amoeba move through deformation of their cell membrane and intercellular movement of their cytoplasm. The presence of plastics may affect protist movement and speed as plastic particles can either block their path, or, if in contact or ingested, become a heavy burden or toxic, potentially altering protist migratory routes and ecological roles. Few studies have shown that soil protists can ingest and excrete plastic particles [13, 14, 15], but yet no research to date has directly isolated or quantified the effects of microplastics on soil protist motility. Soil chips, transparent microfluidic devices containing an artificial soil pore space connectable to natural soil samples, provide a unique opportunity to monitor natural soil community interactions [16, 17]. It is possible to monitor morphological or behavioural changes of soil microorganisms, population- and community dynamics, and interactions of single organisms and communities with their direct environment, including micro-and nanoplastic particles, with the possibility of generating thousands of images and videos. We can monitor morphological and behavioural effects of nanoplastics on single protist cells embedded into their natural communities, but the evaluation of movement trajectories or absolute speed of protists under the microscope is challenging. The rise of automatic AI monitoring technologies has revolutionized ecological research, offering distinct advantages over traditional survey methods. Automation enables the detailed quantification of ecological interactions at unparalleled spatial and temporal resolutions. The rich, multidimensional data collected, from individual behavioral and morphological traits to species abundance and distribution, facilitates the creation of predictive models that integrate data across ecological scales [18]. Our previous work using deep learning method image analysis demonstrates the feasibility to quantify bacteria abundance and gives information about their morpho-diversity: Changes in cellular size and shapes, and the spatial distribution of the cells including cell group formation up to simple biofilms [19].

In this work, we investigated the effects of nanoplastic pollution on soil protists movement using microfluidic soil chips. We hypothesised that the exposure to nanoplastics would change movement trajectories and reduce movement speed of the different protist groups, because of the additional weight of potential ingested particles without nutritional benefit, or potential toxic effects. The soil chips were inoculated with natural soil samples to simulate realistic environmental conditions illustrated in Figure 1 (a). The inner pore spaces of the chips were treated with nanoplastics, where direct interactions and effects could be observed at a cellular level as shown in Figure 1 (b). As depicted in Figure 1 (c-d), to analyze protist behavior, we employed the off-the-shelf convolutional neural network YOLO (You Only Look Once) for tracking and segmenting key protist functional groups, including the morpho-translocational groups amoebae, flagellates, and ciliates. Furthermore, SIFT (Scale-Invariant Feature Transform)[20, 21] was used for feature extraction, and a transformer-based model adapted from LightGlue[22] was developed to comprehensively register protist movements. This approach enabled the quantification of key behavioral metrics, including size, shape, absolute speed, directional changes, and movement trajectories of targeted soil protists. Reduced movement speeds were observed in both flagellates and ciliates at a marginal significance level, while amoebae were unaffected. Visualization revealed that flagellates navigate complex environments by actively sensing their surroundings with flagella, while allocating relatively less energy to propulsion. Overall, this study introduces a novel framework for live tracking and behavioral analysis of soil protists, representing a transformative methodology for investigating soil ecosystems and advancing environmental research, as well as biological and ecological data acquisition at microscale resolution.

**Figure 1:**
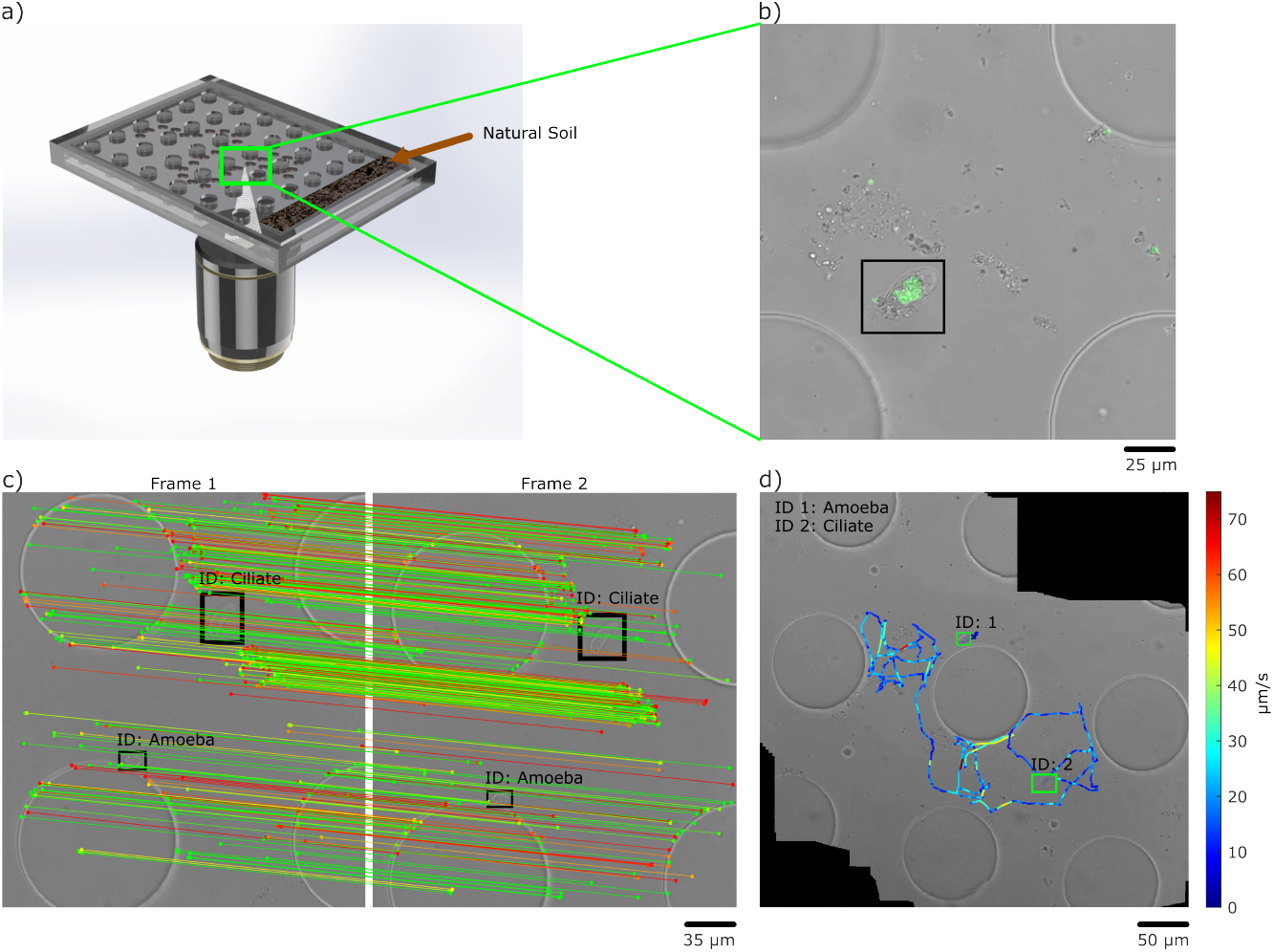
(a) Rendered image of the experimental setup, including a microfluidic ‘soil chip’ and a microscope objective lens. The complete setup includes a microfluidic chip inoculated with a natural soil sample and an inverted microscope for visualizing and tracking protist movements within the chip. (b) Overlay bright-field and epifluorescence images showing the uptake of fluorescent PS nanospheres by a testate amoeba in the soil chip treated with 10 mg/L. (c) Illustration of how video frames of a microscopic video moving across a sample in x-y directions are processed using a transformer-based matching module to track and map individual protists across successive frames. (d) Reconstruction of protist movement trajectories, with absolute movement speeds visualized on the trajectory map using a color scale (in *µ*m/s).

## 2. Experimental methods

### 2.1. Experimental setup

#### 2.1.1. Nanoplastic preparation

The polystyrene (PS) nanospheres used in this study were obtained from Bangs Laboratories Inc. (www.banglabs.com). These nanospheres have a diameter of approximately 60 nm, are carboxylated and negatively charged, and emit fluorescence (excitation: 480 nm, emission: 520 nm) to allow visual monitoring. The preserving solution of these nanospheres contains 0.1% Tween 20 and an antimicrobial compound (2mM sodium azide), components that affect microbial growth and therefore they need to be removed from the solution. To achieve this, dialysis was performed as described by Mafla-Endara et al. [23]. Following this step, Dynamic Light Scattering (DLS) on DynaPro Plate Reader II (Wyatt instruments, USA) was conducted to measure particle size and potential aggregation. The measurement showed no significant changes in particle size. Prior to the incorporation of the nanospheres to the nutrient medium, the nanospheres were coated with bovine serum albumin (BSA) as described by Mafla-Endara et al. [23], to lower self-aggregation and attachment to the PDMS surface.

#### 2.1.2. Microfluidic chips

The microfluidic soil chips serve as observation windows into the soil, featuring a channel height of 10 µm and a regular array of predefined polydimethylsiloxane (PDMS) pillars with a diameter of 100 *µ*m and 75 *µ*m gap between each one. The master for the chip design was fabricated via photolithography using a maskless aligner (MLA150, HEIDELBERG INSTRUMENTS). The pattern was directly written on a 10 *µ*m thick layer of mr-DWL 5 photoresist (MICRO RESIST TECHNOLOGY). PDMS was subsequently cast onto the master to replicate the channel pattern, followed by plasma bonding to a coverslip to create a complete chip. The coverslip with the chip upon was glued into a hand-cut hole in a Petri dish. These soil chips enable the direct observation of soil protists capable of migrating from natural soil samples into the artificial microstructures.

#### 2.1.3. Soil chip set-up

Each chip was filled with 20 *µ*l of a nutrient medium containing 4 *µ*l malt medium, 0.2 *µ*l BSA, and varying volumes of PS nanosphere solutions in ddH2O to obtain a final concentration of 0mg/L, 2mg/L and 10mg/L (n=5, total chips=15). The remaining volume was filled up by ddH2O. Both concentrations of PS nanospheres were chosen for evaluating the behavior of protists under medium and high nanoplastics exposure scenarios, concentrations that have earlier been shown to affect protists [23] and are environmentally relevant. Soil samples (0-10 cm) were collected from a meadow in the vicinities of Lund, Sweden (55.711n; 13.264e) in July 2024. A block of soil (3.25 g) moistened with 1ml of water was placed in contact with the entry of the chip. A sterile wet tissue was placed in the Petri dish and it was sealed with Parafilm to prevent evaporation. To maintain a humid environment, the chips were stored in plastic boxes and left at room temperature. The chips were inoculated on the 4th of July 2024 followed by video acquisition on day 8 and 18 as we have found from previous experiments that diversity peaks around 2 weeks.

#### 2.1.4. Video acquisition

Videos of three classes of soil protists were recorded using a Nikon Ti2-E inverted light microscope equipped with a Nikon Qi2 camera. Video acquisition was performed treatment-blind to prevent observation bias. The videos were captured at 11 frames per second (fps) or 30 fps. The 11 fps videos were taken with a 40X objective and a 1.5X magnification lens, with an analog gain of 1X, an exposure time of 20 µm, and a 14-bit color depth. The condenser was fully open, and the resolution of these videos was 2136 x 2136 pixels, with a spatial scale of 0.12 µm/pixel. For the 30 fps videos, a 100X objective was used, with a 2.8X analog gain, an 11 ms exposure time, 14-bit color depth, and the condenser 50% closed. These videos had a resolution of 712 x 712 pixels and a spatial scale of 0.22 µm/pixel.

### 2.2. Protist tracking

The proposed model comprises two main modules: a detection and tracking module and a Transformer-based matching module. The detection and tracking module detects and tracks the protists’ locations using the trained YOLOv8m model, which generates masks to eliminate dynamic features introduced by the fast movement of protists. These masks are applied to the corresponding image frames, isolating stable regions for further processing. The Transformer-based matching module then matches features across images and performs image registration tasks. The overall architecture of the model is illustrated in Fig. 2.

**Figure 2:**
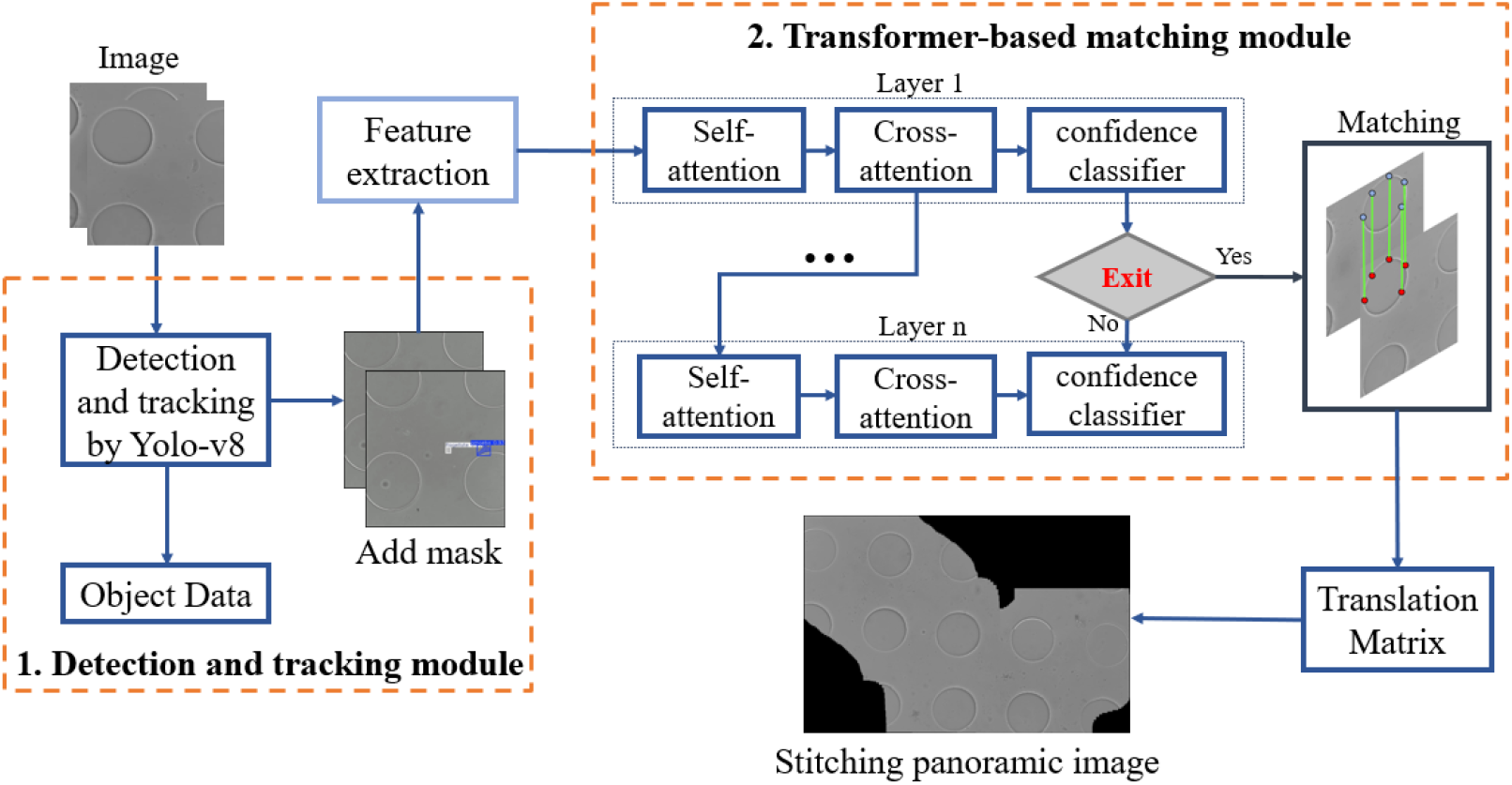
Architecture of the proposed system for stitching panoramic images of soil protist movement. The system comprises two main modules: Detection and Tracking Module and Transformer-based Matching Module. The iterative process continues until the matching confidence threshold is met. Matched features are used to compute a translation matrix, enabling accurate panoramic image stitching.

#### 2.2.1. Protist tracking model training and evaluation

The training datasets were images sampled from video frames of protists in the microfluidic chips. The sampling rate of the video clips varied depending on the type of protist in each video. For example, amoebae, which typically move very slowly, were sampled at a rate of 0.1 frames per second, while ciliates and flagellates, which move more rapidly, were sampled at 1 frame per second. The images were manually annotated using Roboflow [24]. A total of 2,007 images were included in the dataset, which was split into training, validation, and test sets with a ratio of 7:2:1. The model was then trained using supercomputers provided by the National Academic Infrastructure for Supercomputing in Sweden (NAISS) at the Chalmers Centre for Computational Science and Engineering (C3SE). Four Nvidia A40 GPUs were utilized to train the YOLOv8m object detection and tracking models powered by Ultralytics. Performance evaluation was conducted on the test dataset, yielding the following results as shown in table 1. Each frame required only 4 milliseconds for inference, making the model fast enough for future real-time applications.

**Table 1:** Performance metrics of the YOLOv8m model trained for protist tracking.

|  | Precision | Recall | mAP@50 | mAP@50-95 |
| --- | --- | --- | --- | --- |
| Bounding box | 0.942 | 0.929 | 0.952 | 0.848 |
| Masks | 0.953 | 0.953 | 0.967 | 0.775 |

### 2.3. Video compensation

SIFT was employed to extract feature points from the masked images frame by frame. SIFT demonstrates superior performance in sparse feature scenarios, providing stable and accurate feature point extraction. Experiments comparing various extraction methods confirmed that SIFT outperformed alternatives like Superpoint [25], Disk [26] and XFeat [27] for this task. The extracted feature points, along with their visual descriptors and positional data, are fed into the Transformer-based matching module for further processing. To evaluate the performance of feature extraction, we compared the capability of several extractors under microscopy data. We used 560*640 images with each image set to detect up to 1000 feature points. In order to estimate the number of effective feature points, we take the fixed pillar in the microscope image as the effective feature region, count the 15 pixel coordinate points near the circumference of the pillar as the effective feature points, and calculate the evaluated metric value. The evaluated results are shown in Table 2. The feature points extracted by SIFT are more distributed near the pillar, which can well reduce the error of the later matching step. Although Superpoint features are more evenly distributed in the image, it rather elevates the matching error in the alignment task of this study.

**Table 2:** Feature Extraction Methods Comparisons.

|  | Average keypoints | Valid keypoints |
| --- | --- | --- |
| Superpoint | 420 | 14.2% |
| Disk | 1000 | 15.6% |
| XFeat | 419 | 40.7% |
| **SIFT (Best)** | **733** | **75.3%** |

**Table 3:** Matching Model Comparisons.

|  | Match pairs | Valid Matches | Time (ms) |
| --- | --- | --- | --- |
| SIFT + BFM | 287 | 20.8% | 8.2 |
| SIFT + OmniGlue | 120 | 21.1% | 80.3 |
| SIFT + SuperGlue | 71 | 32.3% | 79.6 |
| SIFT + Sgmnet | 41 | 71.2% | 78.6 |
| **SIFT + Our (Best)** | **288** | **83.3%** | **50.1** |

Using the features extracted by SIFT, our method is then compared with BFM (Brute Force Matching), Omniglue [28] and Sgmnet [29], for the evaluation of the number of matches pairs. Meanwhile, considering the similarity and dynamic features of microscope image features, we added the evaluation of effective matching pairs to ensure that the matching pairs can be used for accurate image alignment. As can be seen from the data in the table, our method outperforms BFM, Omniglue and Sgmnet in terms of the number of matches pairs and effective matching pairs on SIFT features, and reduces the inference time by 20%. BFM, as the simplest and fastest 2D feature point matching method, can perform well in traditional image matching tasks. However, in microscope images with a large number of similar features, there are large number of matching errors. Omniglue and Sgmnet are both pre-trained neural network-based matching methods that can cope with the changes in complex environments, but in this task, the sparse similarity of the features greatly reduces the number of matching pairs and stability. Our method outperforms the other methods in both the number of matching pairs and effective matching pairs, and maintains high accuracy and fast matching speed while ensuring matching stability.

#### 2.3.1. Matching by Transformer

In the matching task, the transformer’s attention module allows the model to learn the interrelationships of key points in the images and establish connections between two different images. However, limited by the traditional transformer structure, the model training difficulty grows quadratically with the complexity of the image and the number of keypoints. In this study, the microfluidic images have very few feature points and consist almost entirely of grey and white pixels. The microfluidic features in the region are again very similar. Meanwhile, a fast moving Protist produces a large number of dynamic features. All these problems led to a decrease in the matching accuracy, resulting in biased image registration. Therefore, based on LightGlue [22], a lightweight transformer-based feature matching model that effectively reduces model size while preserving matching accuracy, we have developed an image matching model adapted for protists microscopy image features. The Transformer-based matching module consists of four key components: positional encoding, attention mechanisms, and a matching head.

##### 1. Positional Encoding

Firstly, position encoding is performed, which allows the model to capture the relative spatial relationships between feature points. The 2D image coordinates are normalised to the range [-1, 1] and the aspect ratio of the image is preserved. These coordinates are then mapped into frequency space by a linear projection. This projection transforms the 2D coordinates into a frequency representation suitable for coding. Specifically, for each feature point pair, the model computes a vector of relative positions between them and then rotates the frequencies according to this vector. After encoding, it is input into the attention layer.

##### 2. Attention Mechanisms

The Self-attention unit learns features from one image and each feature point focuses on other feature points to learn contextual information from the same image. Then, the Cross-attention unit performs cross-learning of features from two images. In this way, the model can identify and match feature points across images more accurately.

##### 3. Confidence Classifier

The Confidence classifier unit evaluates the reliability of the model’s predictions at each layer to determine whether additional computation is necessary. At the end of each layer, the Confidence classifier predicts a confidence score for each feature point, determining whether further processing is required. If the confidence scores for most of the feature points are above the threshold and a predefined confidence ratio is reached, the model can be considered to have enough confidence in the matching of these feature points to terminate the computation early. The Confidence classifier enables the model to improve speed without sacrificing accuracy by dynamically adjusting network depth based on the difficulty of each image pair.

##### 4. Matching Head

In the matching unit, the model calculates the similarity scores between all pairs of feature points in the two images to form a matrix of matching scores *S*. Each element of this matrix, *S*_*ij*_, represents the similarity between the *i*th feature point of image A and the *j*th feature point of image B. This matrix is then used to calculate the similarity score between the two images. This score is calculated using the following equation:

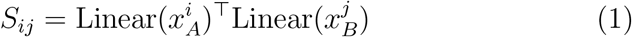

Where 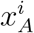 and 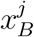 are the updated state vectors of the *i*th feature point in image A and the *j*th feature point in image B, respectively, and *Linear*(·) denotes a linear transformation used to map the state vectors to the similarity score space.

The model also computes a matchability score *σ*_*i*_ for each feature point, indicating whether the feature point is likely to find a match in another image. This score is computed by a sigmoid function:

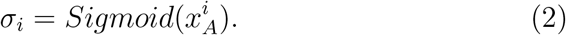

Sigmoid function maps the state vector to the interval [0, 1], indicating the likelihood of a match.

Combining the pairing score and the matchability score, model constructs a soft assignment matrix *P*. This matrix is computed using the following equation:

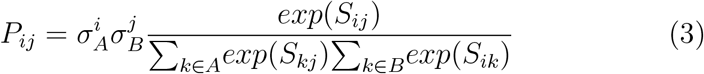

Where, 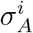 and 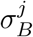 are the matchability scores of the *i*th feature point in image A and the *j*th feature point in image B, respectively. This formula ensures that the sum of matching probabilities for each feature point is 1, thus forming a probability distribution. Finally, from the soft allocation matrix *P* from which the matching pairs are extracted. Feature point matching between two images is predicted by calculating pairing scores and matchability scores between feature points and then combining these scores. This approach allows the model to take into account the geometric and visual similarity of the feature points along with the matchability of the feature points, thus improving the accuracy and robustness of the matching.

### 2.4. Statistical analysis of the protist data

Segmentation masks and positions of the morpho-/locomotion type groups flagellates, ciliates, and amoebae were extracted from the videos and calculated into average size, coefficient of variation of size, maximum speed, average speed, and direction change rate. To mitigate the effects of ID switching, we first examined the frequency distribution of speed across all three functional groups and removed any detection errors defined as values exceeding the 99.5th percentile. Data analysis, statistical computations, and figure plotting were performed using MATLAB. Maximum speed was determined by smoothing the speed data using a span of 3 adjacent individual time points and then identifying the peak value. Subsequently, that information was statistically analyzed with a linear regression model for relationships to the response variables with increasing concentrations of NPs. Data of the linear regression model was checked for normal distribution of their residuals; data with non-normally distributed residuals were log-transformed to meet requirements for the linear regression model.

## 3. Results and discussion

### 3.1. Protist behaviour

In total, 259 videos were taken and processed successfully. 121 flagellates, 44 ciliates, and 94 amoebae were monitored and followed under the microscope, and 1-30 mins long videos recorded, from which the protist tracking was developed. We observed changes in the mobility of soil flagellates and ciliates upon high nanoplastic exposure (Fig. 4 a) and 3), reflected in reduced average and maximum speeds. As shown in Figure 4 a), soil flagellates showed a 24% reduction in average speed at highest concentrations of NPs, compared to the non-polluted control, at a borderline significant relationship (**p** = 0.083) in a linear regression model applied to the log-transformed average speed data. Similarly, a linear regression model was applied to the log-transformed average speed data of the ciliate group. A 28% reduction in average speed and a 30% reduction in max speed was observed compared to the control for the ciliates, also at a marginally significant difference (**p** = 0.056 and p= 0.049). In future experiments, more chips and/or longer observation times should be used to compensate for the fact that ciliates are not occurring as frequently as other protist types in soils [30]. Overall, amoebae exhibit significantly slower movement speeds compared to flagellates and ciliates, although exceptions exist. However, no significant difference in amoeboid movement speed was observed across different nanoplastic treatments.

There are several mechanisms by which protists interact with nanoparticles. Many protists, such as amoebae, ciliates, and flagellates, use phagocytosis to ingest particles, including nanoparticles. [31] Alternatively, nanoparticles can also adhere to the protists’ external membrane due to electrostatic interactions or hydrophobic affinity. Positively charged nanoparticles are particularly prone to binding, given the generally negative surface charge of protist membranes [32]. Surface attachment itself does not require energy expenditure by the protist but can interfere with normal membrane function or motility as the plastic particles constitute extra weight to carry around. Another way is passive diffusion to the intercellular space. As we used nanoparticles at 60 nm in our study, it is unlikely they passively diffuse through protist membranes. In our work, the nanoparticles were seen taken up and transported by the soil protists, as shown in Figure 1 (b) and figures presented by Mafla-Endara [33]. As plastic nanoparticles constitute a weight with no nutritional value to carry, and a potential intracellular uptake can interfere with cellular processes, our hypothesis was to see a significant trend of movement speed reduction as the nanoplastic environmental concentration rises.

Notably, most of the above-named previous studies focus on single specific species, whereas our findings are based on morphotype-level observations of many different protist species in a realistic microbial community. While our dataset remains limited, we anticipate that full automation of monitoring processes will enable the collection of larger datasets, ultimately leading to more accurate and robust conclusions.

In soils, protists facilitate the movement of nanoparticles across microenvironments, such as from bulk soil pore spaces to root zones. This was evident in our experiments where protists carrying nanoparticles on their cellular in- or outside navigated through artificial pore spaces, as shown in the trajectory figures. The trajectory of the flagellate shown in Figure 4 b) highlights the interaction with its surroundings. The color of the trajectory path corresponds to the instantaneous speed. The flagellate navigates through the space, including a fungal hypha and soil particles, at very low speed. However, there is a noticeable surge in speed when it attempts to squeeze through the fungal hypha, showing that it actively sensed its surroundings using its flagella and reacted to obstacles. In contrast, ciliates have the highest average moving speed as shown in Figure 3 a). In comparison to the flagellates, the ciliate trajectory shown in Figure 3 b) reveals many direction changes while moving through the complex environment. Protists are foundational links in many food webs. When they ingest nanoparticles, they can transfer them to higher trophic levels, such as invertebrates, although our soil chip experiments lacked these larger organisms. However, we observed trophic interactions between protists preying on one another, suggesting the potential for nanoparticle transfer even within protist communities. This trophic transfer raises concerns about nanoparticle persistence and potential toxicity as they move up the food chain. In this work, we use a deep learning approach to extract segmentation masks of the target cells, allowing us to analyze changes in their shape. This makes it possible to study oscillatory changes in amoebal cell shape and their relationship to motility. Recent research suggests that amoeboid cells exhibit faster movement when their shape is highly dynamic and undergoes significant changes. Alonso-Latorre et al. highlights that amoeboid cells rely on cyclic shape changes, such as dilation, elongation, and protrusion/retraction, to generate traction forces. Cells with greater shape deformation during their motility cycle exhibit faster migration speeds. [34] Computational models also confirm amoeboid cells with more extensive and rapid shape changes move faster, particularly when these deformations are coordinated with intracellular polarity and substrate interactions. [35] Although we did not observe significant differences in average movement speed across the different nanoplastic treatments, we found a strong correlation between average speed and the coefficient of variation as shown in Figure 5 a) and c). The coefficient of variation in size is defined as the ratio of the standard deviation to the mean size. It quantifies the relative magnitude of size fluctuations in amoebae during locomotion, with a higher value indicating greater variability relative to the mean. Our analysis revealed that amoebae with greater shape deformation during their motility cycle exhibited faster migration speeds.

**Figure 3:**
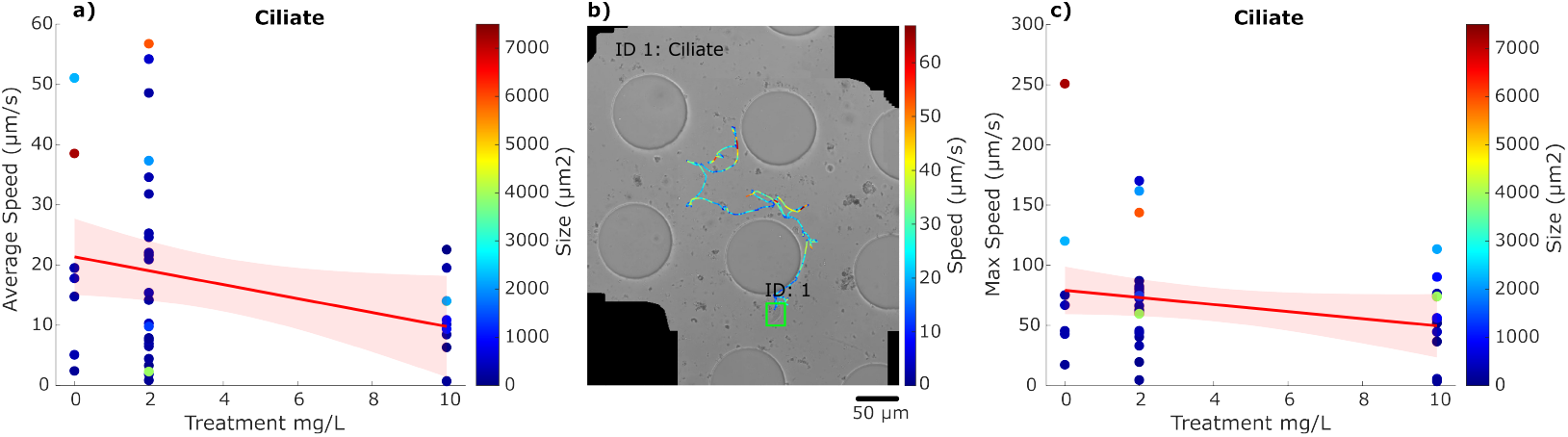
(a) Average movement speeds of individual ciliates under increasing polystyrene nanoplastic concentrations. The color of each marker indicates the size obtained from the corresponding ciliate segmentation. The red line indicates a linear regression with 95% confidence interval. (b) The trajectory of an individual ciliate in a complex environment is shown, with the color along the path representing the instantaneous velocity at each point. c) Maximum movement speeds of individual ciliates under increasing polystyrene nanoplastic concentrations. The color of each marker corresponds to the size obtained from the corresponding ciliate segmentation. The red line indicates a linear regression with 95% confidence interval.

**Figure 4:**
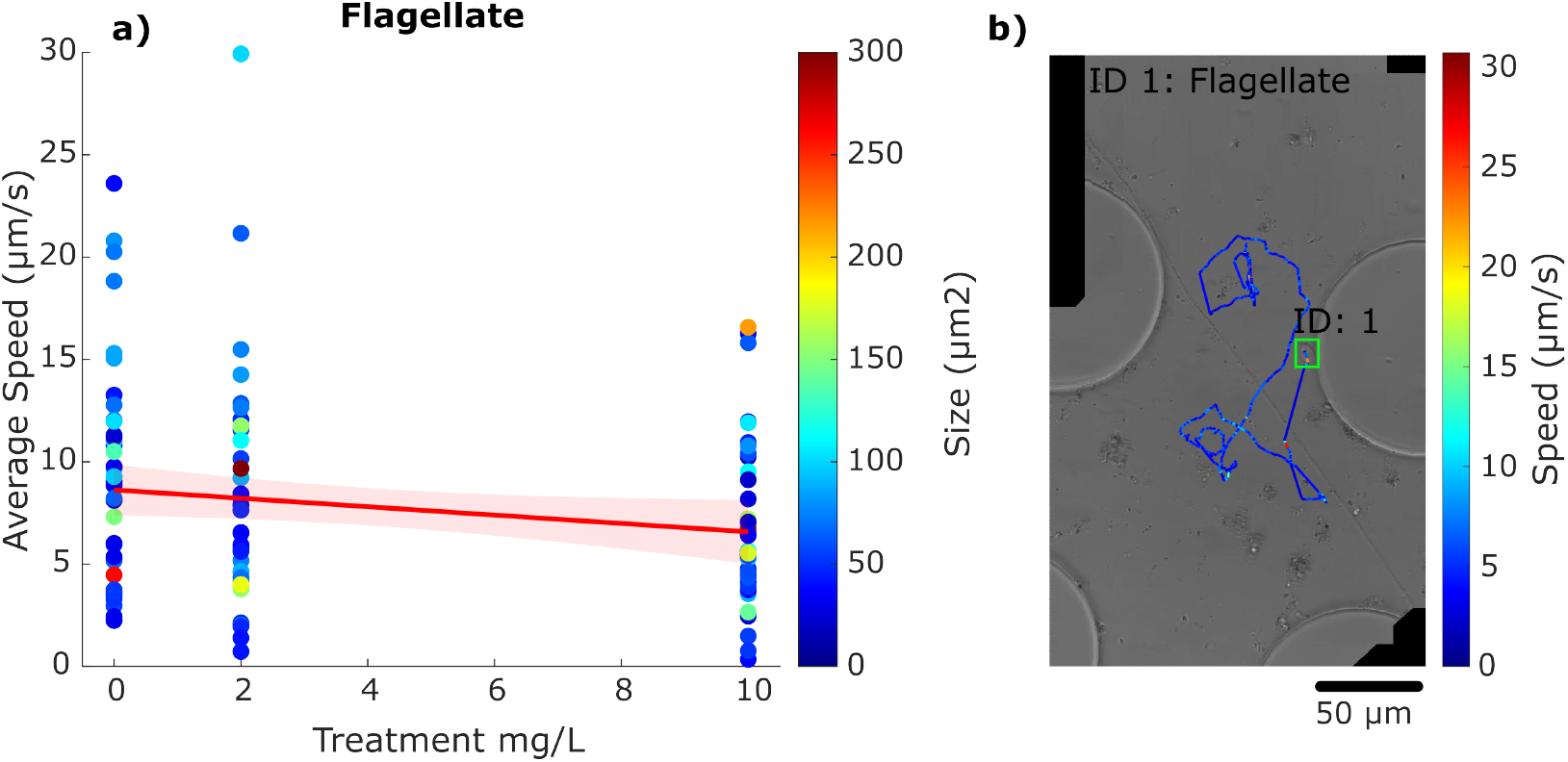
(a) Average movement speed of individual flagellates under different polystyrene nanoplastic concentrations. The red line indicates a linear regression with 95% confidence interval. (b) Individual trajectory of a flaggelate in response to different environmental stimuli. The color and size of each marker represent the size of the corresponding flagellate segmentation.

**Figure 5:**
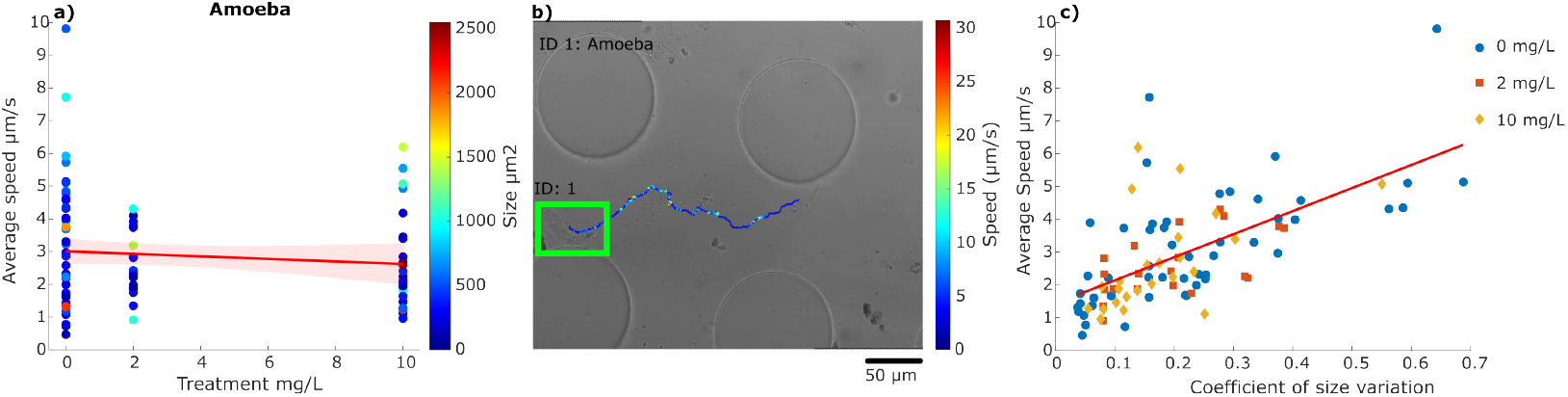
(a) Average movement speeds of individual amoebae plotted against increasing polystyrene nanoplastic concentrations. Colour labels represent their average size. The red line indicates a linear regression with 95% confidence interval. (b) Individual trajectory of an amoeba moving towards the left side of the field of view. Trajection color marks the instantaneous velocity at each point. (c) Average move ment speed of individual amoebae plotted against their coefficiant of vairation in size, with a higher coefficient indicating greater relative variability during movement. Marker color and shape correspond to different nanoplastic treatments: blue for 0 mg/L, orange for 2 mg/L, and yellow for 10 mg/L.

Our previous work demonstrated that combining soil chips with AI-driven image analysis enables direct investigation of bacterial communities in soil [19]. Here, we extended the idea to protists, their predators. Many soil protists are highly motile and serve as key vectors in the transport of plastic particles, as shown here, but the impact of the distribution of plastic contaminations into different soil pore spaces remains completely unknown. Our approach opens up novel ways of direct investigation of food web interactions and their impacts on soil processes. This is critical because the protists’ mobility and ecological versatility make them key players in the environmental transport of nanoparticles.

For the first time, the method presented in this work enables visualization and quantification of natural movement trajectories of soil protists under nanoplastic stress, as well as their responses to environmental factors such as soil particles and structural features like pillars. This monitoring system allows us to observe soil protists in a manner similar to studying animals, prompting us to consider how these organisms interact with their surrounding environment. Additionally, we can collect statistical data on their movement speed and variation in velocity and direction in response to various parameters, providing valuable insights into their behavior and adaptation. The data shown in this study illustrate the power of automatic monitoring and its future application in tracking microbial responses to various chemical and physical soil hazards. han

## 4. Conclusions

Our interdisciplinary approach integrates an AI-based tracking system, transformer networks for video frame interpolation to accurately reconstruct protist trajectories and movement speeds. The use of soil chips makes it possible to perform those studies on soil dwelling protists which has previously been impossible for most taxa, as it allows the organisms to move from an undisturbed soil sample into the transparent chip volume without damage that an extraction would cause. The results reveal a reduction in the mobility speed of both ciliates and flagellates as a response to environmentally relevant nanoplastic concentrations, and their active transport of those. Ultimately, our platform marks a major advance toward automated behavioral analysis of microorganisms and can be readily adapted to wide application of microorganisms responses to a range of chemical or physical soil hazards.

## Acknowledgments

The AI training was enabled by resources provided by the National Academic Infrastructure for Supercomputing in Sweden (NAISS), partially funded by the Swedish Research Council through grant agreement no. 2022-06725. We acknowledge financial support by NanoLund. The authors extend their gratitude to Prof. Bo Bernhardsson from the Department of Automatic Control, Lund University for co-supervising master’s students Zuoyi Yu and Jingmo Bai for their valuable preliminary contributions through their master’s thesis work. We acknowledge Familjen Kamprads stiftelse (No. 20230084) to Hanbang Zou. This work is supported by a Future Research Leader grant from the Swedish Foundation for Strategic Research SSF18-0089 to E.Hammer and by BECC, the strategic research environment for Biodiversity and Ecosystem services in a Changing Climate.

